# Interpretable Machine Learning Reveals Complementary Age-Related Signatures in the Oral and Gut Microbiome

**DOI:** 10.64898/2026.09.09.750358

**Authors:** Chowdhury Aseer Ruthbah, Talim Hossain Sadi, Nur E Shiratun Jahan, Abu Nayem Md. Tanzim Adib

**Author notes:** **Corresponding author:** Chowdhury Aseer Ruthbah (;). These authors contributed equally as co-first authors (Chowdhury Aseer Ruthbah, Talim Hossain Sadi). These authors contributed equally as co-second authors (Nur E Shiratun Jahan, Abu Nayem Md. Tanzim Adib).

## Abstract

Whether combining microbiome data from multiple body sites improves prediction, and whether different sites carry complementary or redundant information, are distinct questions that most studies conflate into a single accuracy metric. This work makes two contributions, one methodological and one biological, using paired stool and oral cavity microbiome samples from two independent cohorts. In the primary cohort (Ferretti et al., 2018; n=44), we show that a subject-matched fusion design combined with SHAP-based (SHapley Additive exPlanations) site attribution can detect complementary information between body sites even when no measurable accuracy gain results. Gut (stool) composition alone achieved near-perfect classification (area under the receiver operating characteristic curve, AUC = 1.00) of adult versus newborn age-group status, and combined stool-oral models never exceeded this ceiling. Despite the flat accuracy curve, SHAP analysis of the fused model showed that oral cavity features carried 58.1% of total feature importance versus 41.9% for stool, indicating real, non-redundant reliance on both sites. We replicated this finding in an independent, larger cohort (Brito et al., 2016; n=140) using a harder classification task (sex-based classification), where oral cavity again dominated the fused model’s SHAP importance (60.7% versus 39.3% for stool), and a permutation test confirmed the asymmetry was statistically significant (p < 0.005). The site-attribution asymmetry in the primary cohort was directionally consistent but did not reach significance, consistent with limited statistical power at n=44. The taxa driving these patterns include *Malassezia restricta, Staphylococcus epidermidis*, and *Prevotella melaninogenica*, all of which behave in a manner consistent with their established roles as early colonizers of the neonatal gut, skin, and oral cavity, once verified directly against abundance data. These results support a model of oral-gut microbiome maturation as two distinct, complementary processes, and demonstrate that detecting this relationship requires examining a model’s internal reasoning rather than its accuracy alone. The site-attribution method is released as a standalone, reusable Python tool.

## Introduction

The human body hosts distinct microbial communities across different anatomical sites, the gut, the mouth, the skin, and others, each shaped by its own local physiology, environmental exposure, and immune conditions. Two related but distinct questions follow. The first is biological: as a person’s microbiome develops or ages, do these communities change together, tracking a shared underlying process, or do they mature along separate timelines? The second is methodological: if machine learning is used to answer the first question, is a simple accuracy comparison sufficient to determine whether two body sites carry the same information or genuinely different information?

These two questions are closely linked, and answering one well requires taking the other seriously. Most microbiome-based prediction studies focus on a single body site, usually the gut, and report a single accuracy value as evidence of what that site’s microbiome reveals about a person’s age, health, or development. A smaller set of studies has examined more than one site. Huang et al. (2020) showed that skin, oral, and gut microbiomes can each independently predict chronological age in adults, with skin performing best and gut worst. That comparison, however, used separate cohorts rather than the same individuals sampled at multiple sites. More recently, Myers et al. (2025) examined paired oral and gut samples from 242 people directly and found only a moderate correlation between the two sites’ age-prediction patterns. This suggests real, non-overlapping biological signal, consistent with the fact that most microbial species present in one habitat are absent from the other.

This raises a methodological problem. Suppose two body sites are combined into a single predictive model, and the combined model performs no better than the best single site alone. The natural conclusion, drawn in most such studies, is that fusion did not help. This conclusion, however, holds only if accuracy is the sole measure considered. If one site’s signal is already sufficient to solve the task, a fused model has no room to show an accuracy improvement, regardless of whether it continues to rely on the second site’s information internally. Distinguishing these two scenarios, a genuinely redundant second site from a genuinely complementary one that was simply not needed to reach the accuracy ceiling, requires examining the model itself. SHAP (SHapley Additive exPlanations) attributes a model’s predictions to individual input features, and makes this possible. In a fused model, SHAP values can be grouped by the body site each feature originated from, revealing how much the model actually relies on each site regardless of whether that reliance is reflected in the accuracy score.

We test this idea using two independent public datasets, a primary cohort with 44 matched stool-oral subjects (Ferretti et al., 2018) and a replication cohort with 140 matched stool-oral subjects (Brito et al., 2016). For each cohort, we train classifiers on stool-only, oral-only, and early-fusion stool+oral feature sets, apply SHAP site-attribution to the fused model, and test whether the observed site-contribution asymmetry is statistically distinguishable from random feature-to-site reassignment via a permutation test. We further validate that the taxa driving the model’s predictions behave as established early-life microbiology predicts, by checking their per-subject abundance against their SHAP contributions directly. The result is a methodological framework for detecting complementary multi-site signal that survives an accuracy ceiling, validated across two independent cohorts, along with a biological account of how gut and oral microbiomes mature relative to each other in early life. The full analysis pipeline and a reusable implementation of the site-attribution method are publicly available.

### Literature Review

Microbiome-based age and health prediction has developed largely along single-site lines. Gut studies remain the most common, spanning disease classification (inflammatory bowel disease, colorectal cancer, type 2 diabetes) and continuous biological aging clock models. Deep learning approaches, including autoencoders, convolutional-recurrent hybrids, and phylogeny-aware networks, have progressively been layered onto single-site gut pipelines, generally with modest accuracy gains over classical models such as Random Forest and gradient boosting.

Comparatively fewer studies evaluate prediction across more than one body site. Huang et al. (2020) pooled multiple independent, non-subject-matched cohorts to compare age prediction from skin, oral, and gut microbiomes in adults, finding skin the most predictive and gut the least. Myers et al. (2025) benchmarked a transformer-based dimensionality reduction approach across all three sites at larger scale and also examined subject-matched oral-stool pairs directly, finding only moderate cross-site correlation in age-prediction error, consistent with the compositional distinctness of the two habitats. Separately, several groups have pursued multi-view gut microbiome aging clocks (Chen et al., 2022; Wang et al., 2024) that combine multiple representations of a single site, taxonomic and functional pathway profiles, rather than genuinely paired samples from distinct anatomical sites.

Across this body of work, interpretability is typically reported as a secondary output, a feature-importance ranking or SHAP summary for a single body site, rather than used as a primary tool for characterizing how a fused, multi-site model reaches its predictions. We are not aware of prior work that constructs subject-matched, multi-site fusion features, applies SHAP attribution within a single trained fused model to quantify each site’s proportional contribution, and tests the statistical significance of the resulting site-contribution asymmetry via permutation across two independent cohorts. This gap shapes the present study.

## Methods

### Datasets and study design

Primary cohort (Ferretti et al., 2018): We used publicly available shotgun metagenomic sequencing data from this mother-infant microbiome cohort, accessed via the curatedMetagenomicData Bioconductor package (v3.20.0), which provides uniformly processed taxonomic abundance profiles (species-level relative abundances, generated with MetaPhlAn3) alongside curated subject metadata. The cohort collected samples from multiple body sites from the same individuals across two age groups: adults and newborns. We restricted analysis to subjects with matched stool and oral cavity samples, yielding 44 subjects (21 adults, 23 newborns). Skin samples were excluded since all 15 available skin samples were from adults, making a three-site matched analysis infeasible. Classification target: adult vs. newborn (binary).

Replication cohort (Brito et al., 2016): To test whether the site-attribution pattern generalizes beyond a single cohort and a single classification task, we applied the same pipeline to a second independent dataset from the same Bioconductor package. This study collected matched stool and oral cavity samples from 172 subjects, of whom 140 had samples at both sites. Classification target: sex (female vs. male, 74 vs. 66 subjects), deliberately chosen as a harder, more biologically subtle task than adult-vs-newborn to test whether the method’s findings hold when the accuracy ceiling is not trivially reached. For both cohorts, subjects with multiple samples per site were restricted to the first available sample per site to avoid pseudo-replication.

### Preprocessing

For both cohorts, features began from a shared 667-taxon MetaPhlAn3 reference space. Low-variance taxa (variance below 1×10^−4^) were removed per site to reduce noise, then a centered log-ratio (CLR) transformation was applied with a pseudocount of 1×10^−6^ to handle zero abundances. This is standard practice for compositional microbiome data. After filtering, the Ferretti cohort retained 291 stool and 153 oral taxa; the Brito cohort retained 227 stool and 269 oral taxa (Table 1).

**Table 1.** Preprocessing summary by cohort and body site.

| Cohort | Site | Taxa (before / after filtering) | Matched subjects | Classification target |
| --- | --- | --- | --- | --- |
| Ferretti et al. (2018) | Stool | 667 / 291 | 44 | Adult vs. newborn |
| Ferretti et al. (2018) | Oral cavity | 667 / 153 | 44 | Adult vs. newborn |
| Brito et al. (2016) | Stool | 667 / 227 | 140 | Female vs. male |
| Brito et al. (2016) | Oral cavity | 667 / 269 | 140 | Female vs. male |

### Experimental conditions and classification models

For each cohort, we defined three feature-set conditions: stool only, oral cavity only, and an early-fusion combination of both (feature names prefixed by site of origin). For each condition, we trained three classifiers: L1-regularized logistic regression, Random Forest (300 trees), and XGBoost (200 estimators, max depth 3). Performance was estimated using 5-fold stratified cross-validation. Performance was measured as AUC, reported as mean and standard deviation across folds. To confirm that AUC values reflected genuine signal in the primary cohort, we conducted three robustness checks on the best-performing condition (stool, Random Forest): a label-shuffling null baseline (20 permutations); bootstrap 95% confidence intervals on AUC (200 resamples); and a preprocessing sensitivity check under a 10x larger CLR pseudocount and stricter variance filter.

### Interpretability and site-attribution analysis

We applied SHAP (TreeExplainer for Random Forest and XGBoost, LinearExplainer for logistic regression) to identify which taxa drove predictions in each cohort’s best-performing condition. For the fused condition in each cohort, we additionally summed absolute SHAP contributions by site of origin to quantify how much predictive weight came from each body site within a single trained model. This site-attribution step is the paper’s central methodological contribution.

To test whether the observed site-contribution asymmetry was statistically distinguishable from random feature-to-site assignment, we ran a permutation test for each cohort. In each permutation, we randomly reassigned features to sites (preserving site sizes but scrambling which specific columns belonged to each site), recomputed the max-site SHAP share, and built a null distribution across 200 permutations. The p-value is the proportion of null permutations producing a max-site share equal to or greater than the observed value.

For three top-ranked taxa whose age-direction was ambiguous from literature alone (Malassezia restricta, Staphylococcus epidermidis, Prevotella melaninogenica), we generated SHAP dependence plots and computed Pearson correlations between per-subject abundance and SHAP contribution to resolve directionality empirically. The site-attribution method is implemented as a standalone Python tool (microbiome_site_attribution.py) released alongside this paper at https://github.com/Aseer47/oral-gut-microbiome-fusion. All analyses were conducted in Python using scikit-learn, xgboost, and shap; data extraction used R via the curatedMetagenomicData Bioconductor package.

## Results

### The accuracy ceiling: gut microbiome alone separates adults from newborns

All three conditions in the primary cohort achieved high cross-validated AUC (0.96-1.00) across all three model types (Table 2, Figure 1). The best overall result was Random Forest on stool alone (AUC = 1.00 ± 0.00), matched but never exceeded by the fused condition.

**Table 2.** Cross-validated classification performance by body-site condition and model (primary cohort, Ferretti et al., 2018).

| Condition | Model | Mean AUC | Std AUC | # Features |
| --- | --- | --- | --- | --- |
| Stool | Logistic Regression (L1) | 0.990 | 0.020 | 291 |
| Stool | Random Forest | 1.000 | 0.000 | 291 |
| Stool | XGBoost | 0.960 | 0.049 | 291 |
| Oral cavity | Logistic Regression (L1) | 0.970 | 0.040 | 153 |
| Oral cavity | Random Forest | 0.975 | 0.050 | 153 |
| Oral cavity | XGBoost | 0.975 | 0.050 | 153 |
| Stool + Oral cavity | Logistic Regression (L1) | 1.000 | 0.000 | 444 |
| Stool + Oral cavity | Random Forest | 1.000 | 0.000 | 444 |
| Stool + Oral cavity | XGBoost | 0.960 | 0.049 | 444 |

**Figure 1.**
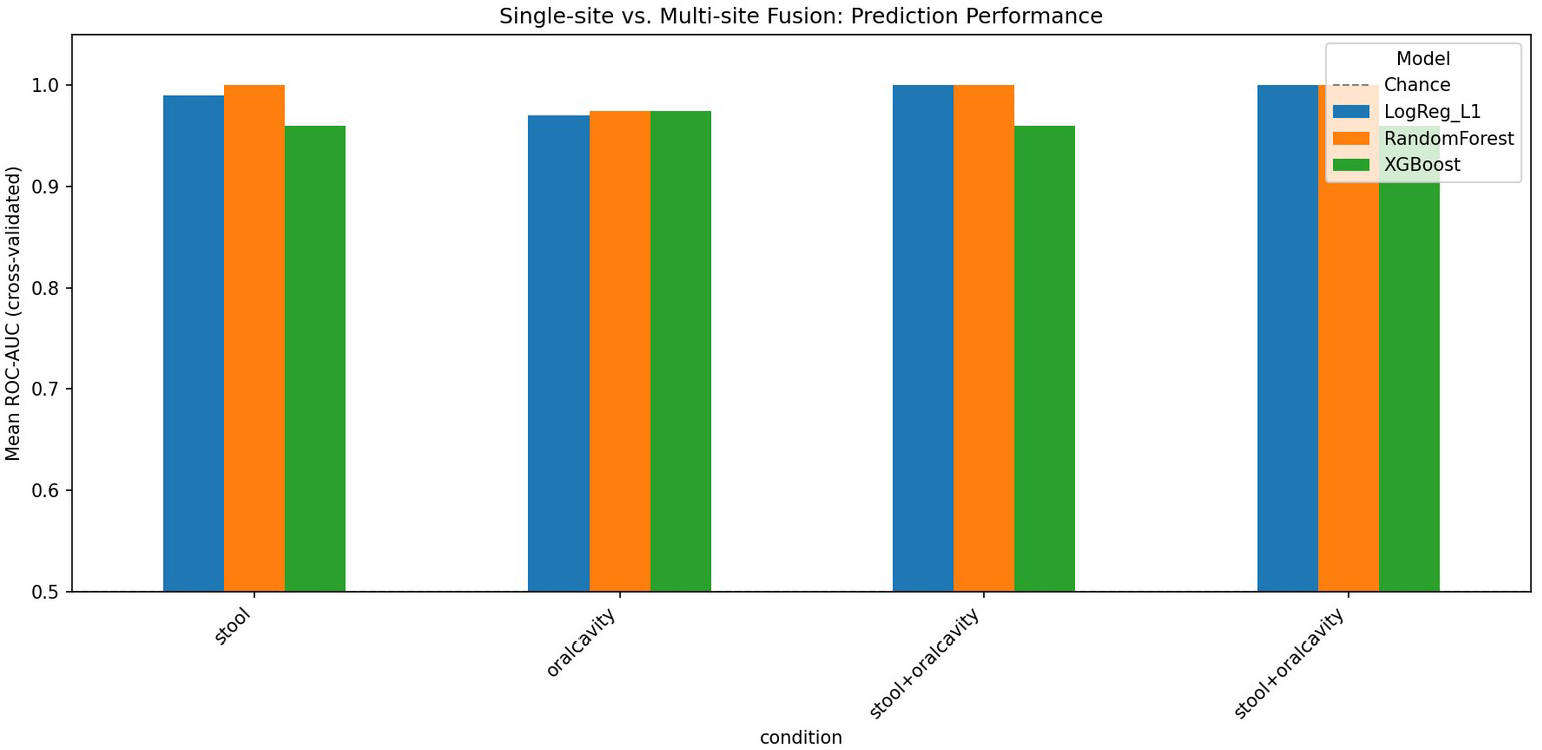
Cross-validated ROC-AUC by body-site condition and model (primary cohort). All conditions cluster near the performance ceiling, with the fused condition matching but not exceeding the best single-site result.

We confirmed the ceiling reflects genuine signal. A label-shuffling null baseline collapsed to chance (mean AUC = 0.500, std = 0.156), ruling out leakage. Bootstrap 95% confidence intervals were narrow and consistently high (stool: 0.997 [0.976, 1.000]; fused: 0.998 [0.982, 1.000]). AUC remained at 1.000 under alternative preprocessing choices (Table 3), indicating the ceiling is a property of the underlying biological signal.

**Table 3.**
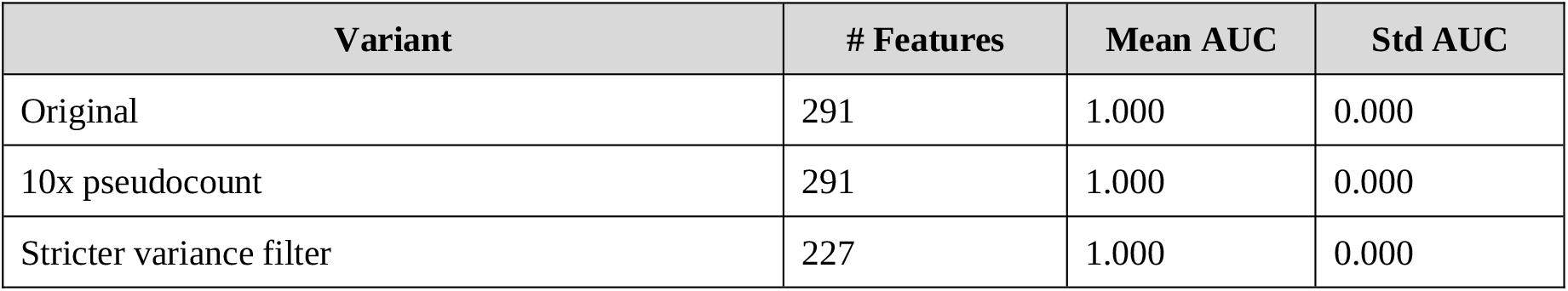
Stool-only Random Forest AUC across preprocessing variants (primary cohort).

| Variant | # Features | Mean AUC | Std AUC |
| --- | --- | --- | --- |
| Original | 291 | 1.000 | 0.000 |
| 10x pseudocount | 291 | 1.000 | 0.000 |
| Stricter variance filter | 227 | 1.000 | 0.000 |

### The methodological finding: the fused model relies on oral cavity information the accuracy score never reveals

Because gut data alone already reaches the accuracy ceiling, ordinary accuracy comparison would conclude that adding oral cavity data is unnecessary. SHAP site-attribution on the fused Random Forest model tells a different story: oral cavity features accounted for 58.1% of total absolute SHAP importance versus 41.9% for stool (Figure 2). The model relies more heavily on oral-derived taxa than on stool-derived ones, despite stool alone being sufficient to reach the same peak accuracy.

**Figure 2.**
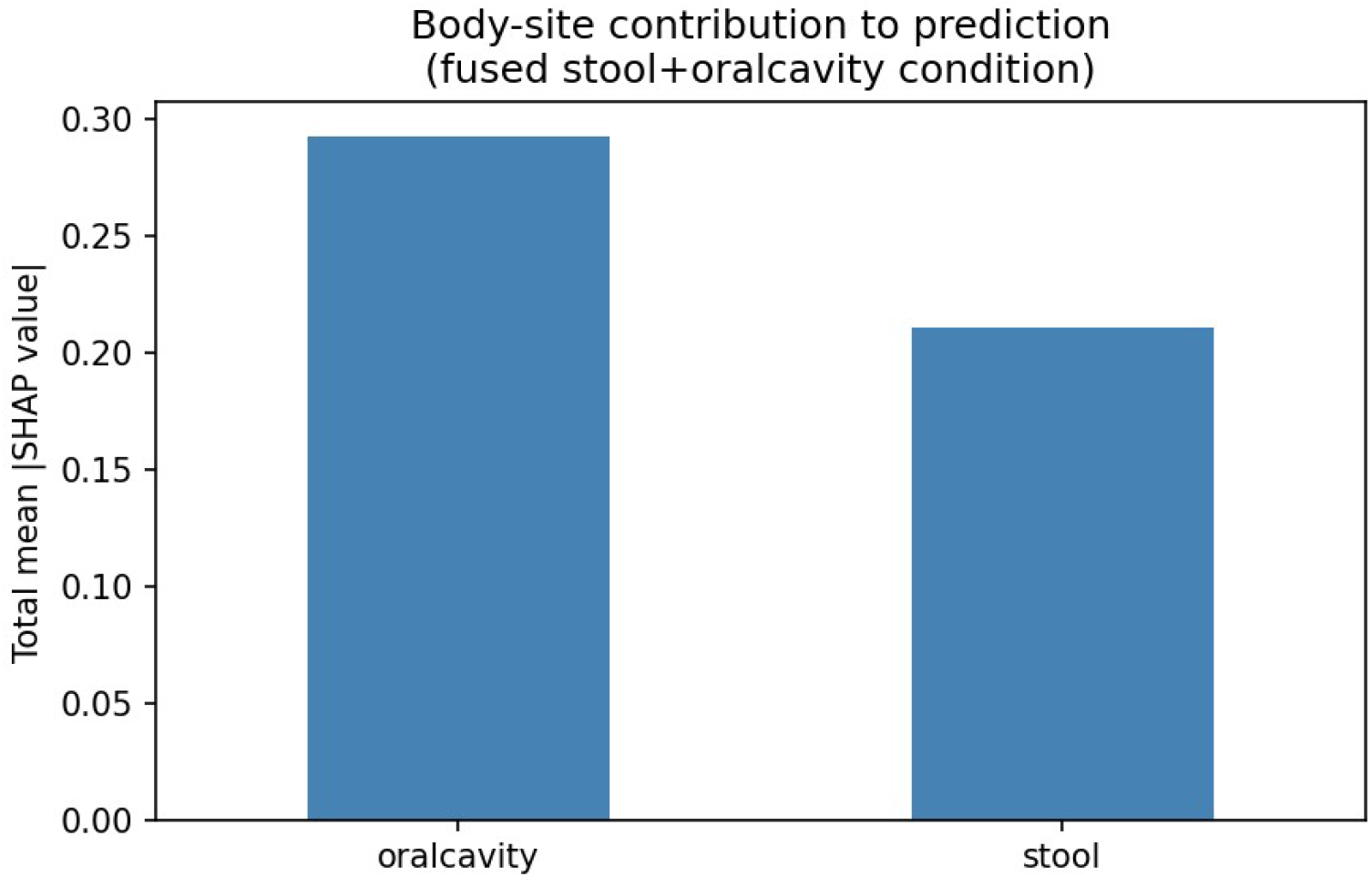
Total mean absolute SHAP contribution by body site of origin in the fused model (primary cohort). Oral cavity features contribute a larger share of model output than stool features, despite stool alone matching the same peak AUC.

The fused-model SHAP summary plot (Figure 3) supports this: oral cavity-derived taxa, including *Escherichia coli, Caulobacter* sp. FWC26, *Prevotella colorans*, and *Prevotella timonensis*, rank among the most influential features alongside stool-derived taxa such as *Malassezia restricta* and *Staphylococcus epidermidis*.

**Figure 3.**
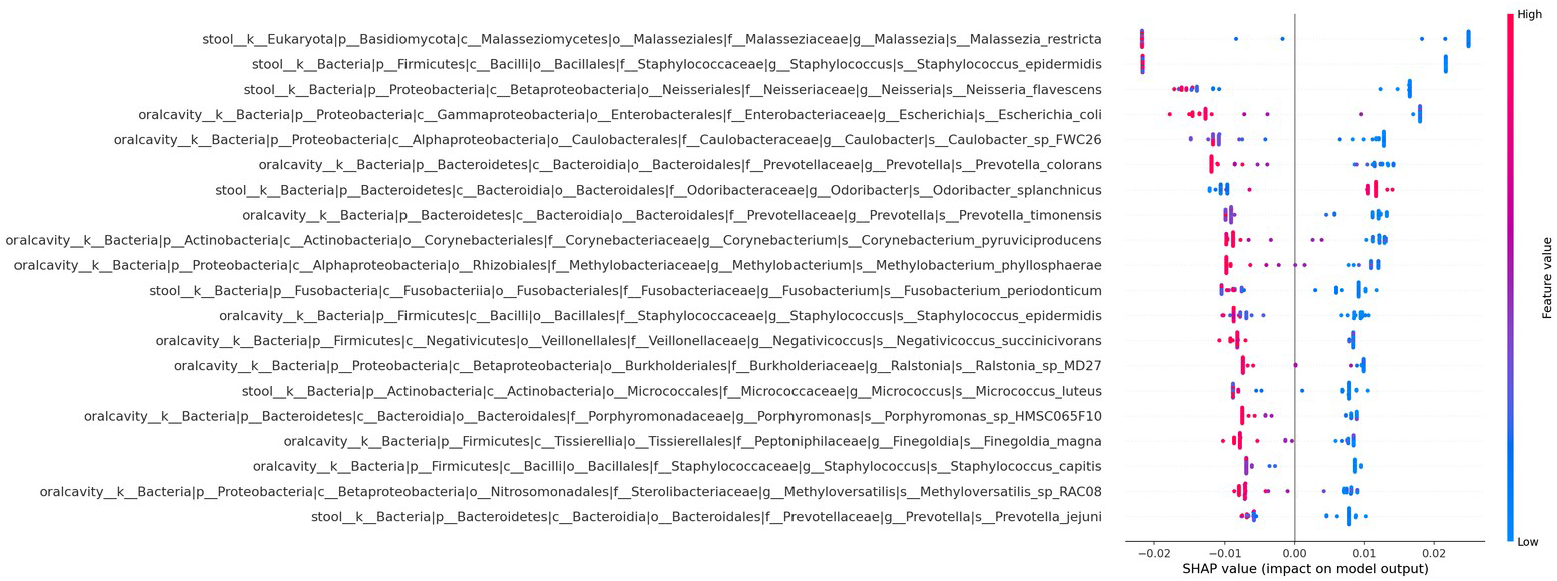
SHAP summary plot for the fused model (primary cohort), feature names prefixed by site of origin.

Permutation test (primary cohort): The observed 58.1% oral share fell within the null distribution (p = 0.89), meaning the asymmetry in the primary cohort alone is not statistically significant. This is consistent with limited statistical power at n=44 and directly motivates the replication analysis below.

### Replication cohort: the site-attribution asymmetry is statistically significant at larger sample size

We applied the identical pipeline to the independent Brito et al. (2016) cohort (n=140 matched subjects) using sex as the classification target, a harder and more biologically subtle task than adult-vs-newborn.

**Table 4.**
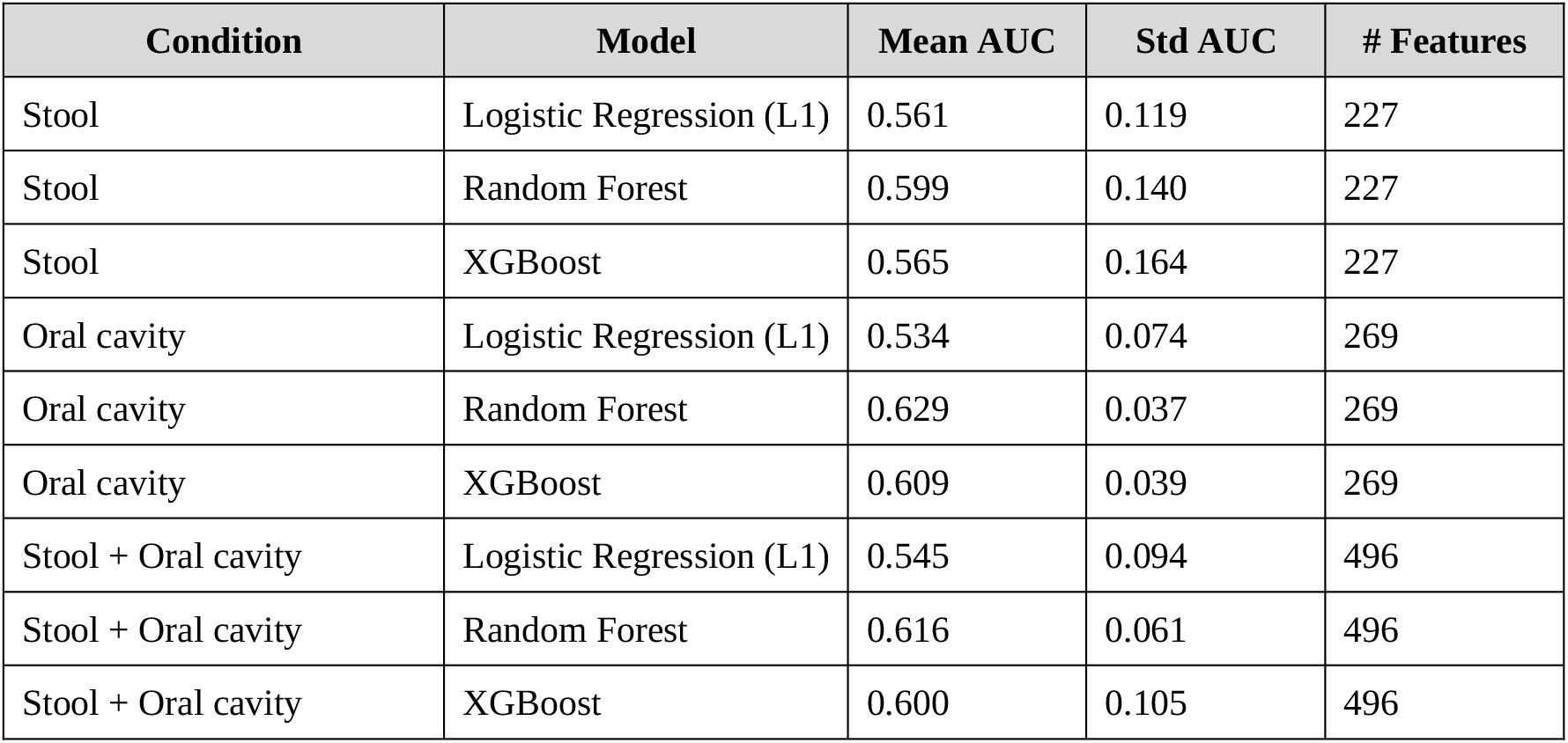
Cross-validated classification performance by body-site condition and model (replication cohort, Brito et al., 2016).

Three findings from the replication cohort are notable. First, the task is genuinely harder: all AUC values are modest (0.53-0.63). Second, oral cavity outperforms stool as the single best site (best oral AUC = 0.629 vs. best stool AUC = 0.599), the reverse of the primary cohort. Third, fusion again fails to exceed the best single-site result.

SHAP site-attribution on the Brito fused model shows oral cavity features contributing 60.7% of total SHAP importance versus 39.3% for stool (Figure 4), closely mirroring the primary cohort (58.1%/41.9%) despite the different cohort, different task, and different accuracy-dominant site.

**Figure 4.**
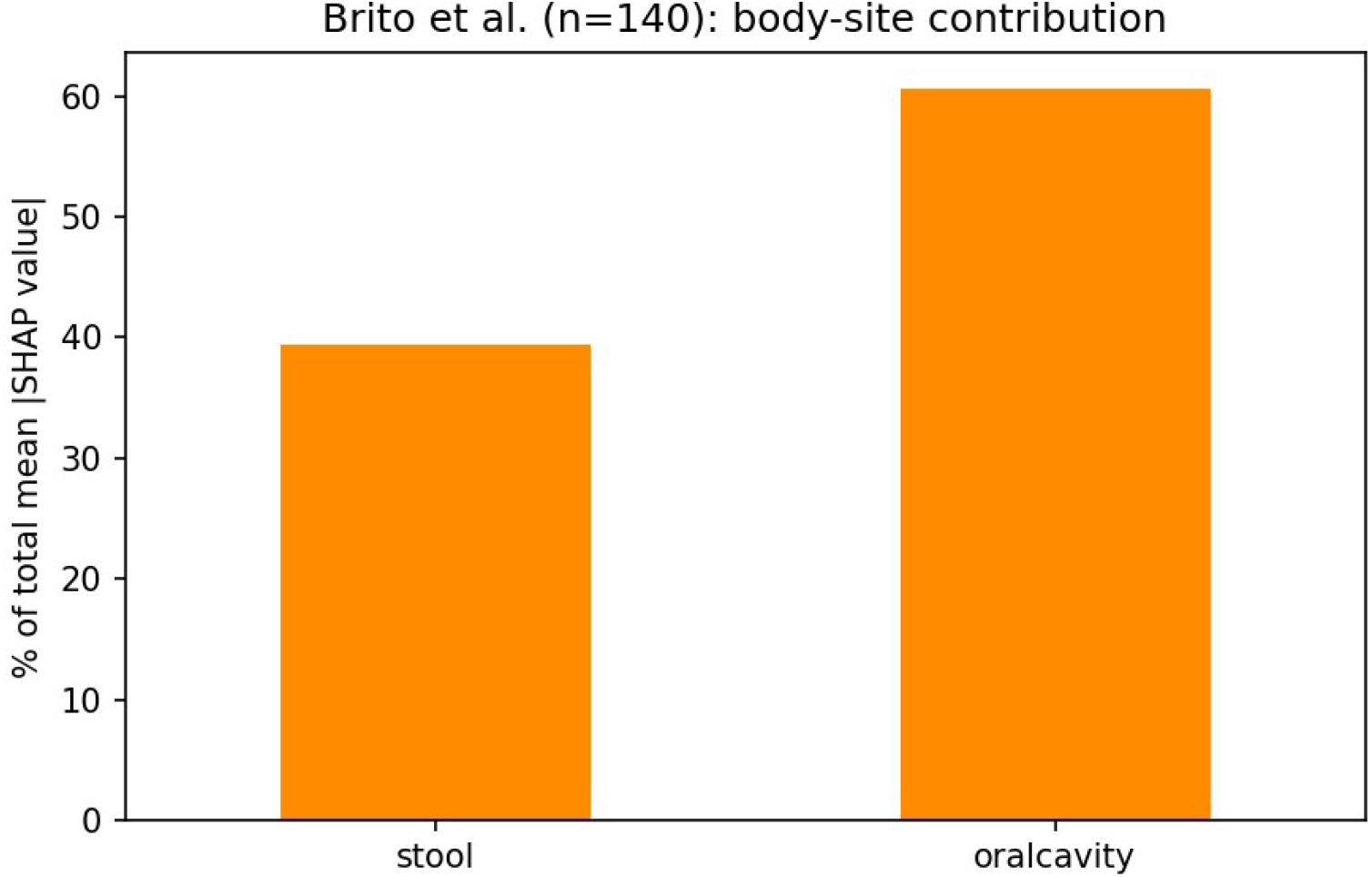
Total mean absolute SHAP contribution by body site of origin in the fused model (replication cohort, n=140). Oral cavity features again contribute more than stool features, closely mirroring the primary cohort result.

Permutation test (replication cohort): With 140 subjects, the observed 60.7% oral share was greater than all 200 permuted null values (p < 0.005), meaning the site-attribution asymmetry in the Brito cohort is statistically significant (Figure 5).

**Figure 5.**
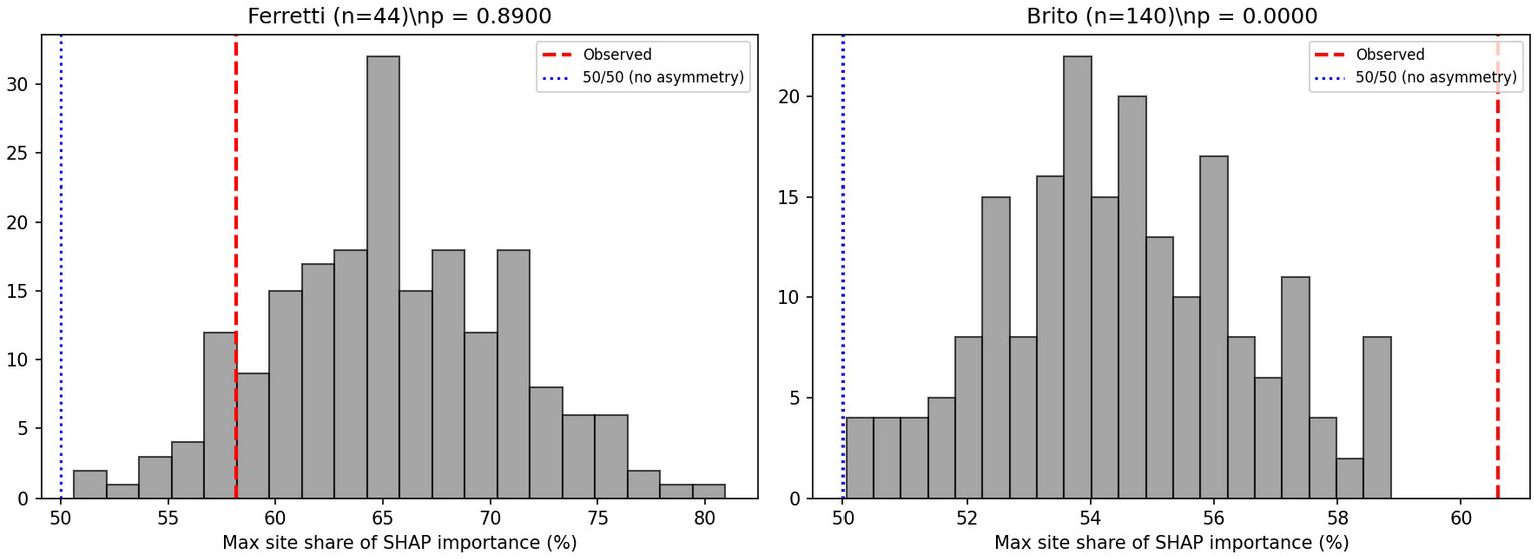
Permutation null distributions for both cohorts. Red dashed line: observed max-site SHAP share. Blue dotted line: 50% (no asymmetry). Ferretti (n=44): p = 0.89, consistent with limited power. Brito (n=140): p < 0.005, statistically significant.

Taken together, these results confirm that the oral-gut site-attribution asymmetry is a real, replicable signal. Its non-significance in the primary cohort reflects insufficient statistical power at n=44 rather than absence of the effect, and its significance in the larger independent cohort confirms the pattern is not a Ferretti-specific artifact.

### The biological finding: key taxa track early-life colonization, confirmed directly

Interpretability analysis on the stool-only Ferretti model (Figure 6) identified *Malassezia restricta, Staphylococcus epidermidis, Neisseria flavescens, Prevotella melaninogenica*, and related taxa as the strongest drivers of the adult/newborn distinction.

**Figure 6.**
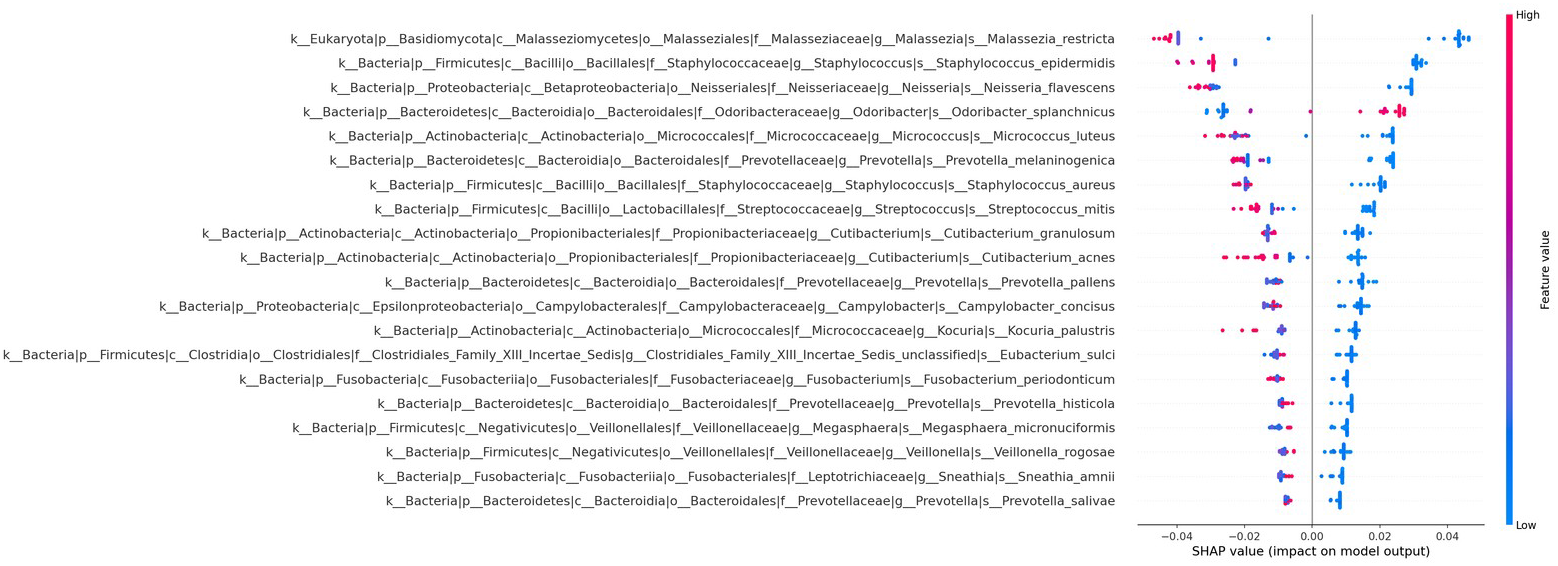
SHAP summary plot for the stool-only Random Forest model (primary cohort), ranking the top 20 taxa by mean absolute SHAP value.

We resolved directional ambiguity by examining each taxon’s actual abundance-SHAP relationship directly (Figure 7). *Malassezia restricta* abundance was strongly negatively correlated with the adult-direction SHAP value (r = -0.74), consistent with longitudinal cohort findings that *Malassezia* abundance decreases over an infant’s first 18 months of life (Hoskinson et al., 2026). *Staphylococcus epidermidis* showed the strongest relationship (r = -0.88), matching its well-documented role as one of the earliest bacterial colonizers of neonatal skin and gut (Datta et al., 2021). *Prevotella melaninogenica* showed the same pattern (r = -0.73), consistent with its established role as one of the earliest bacterial colonizers of the infant oral cavity (Könönen et al., 2022).

**Figure 7.**
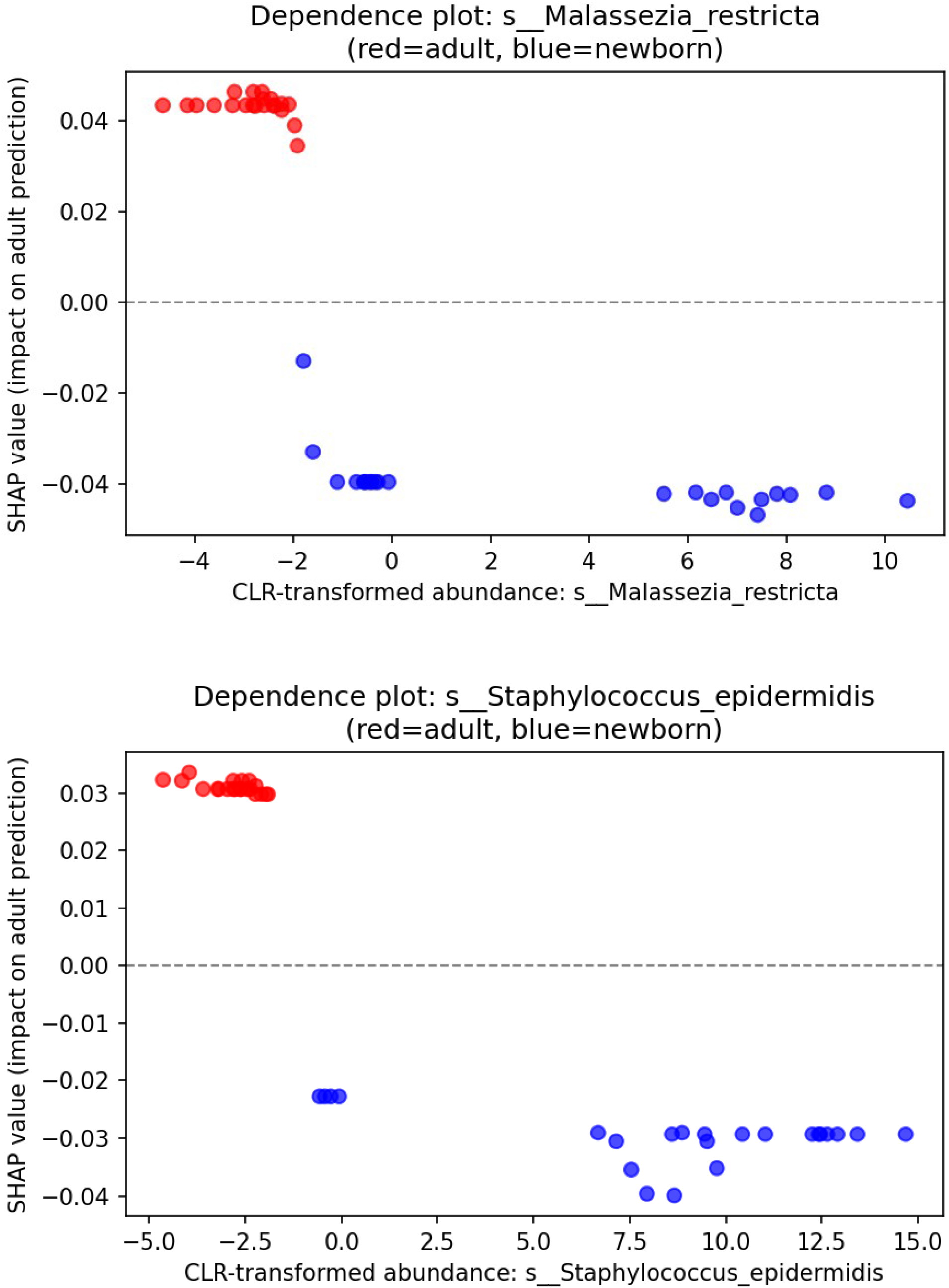
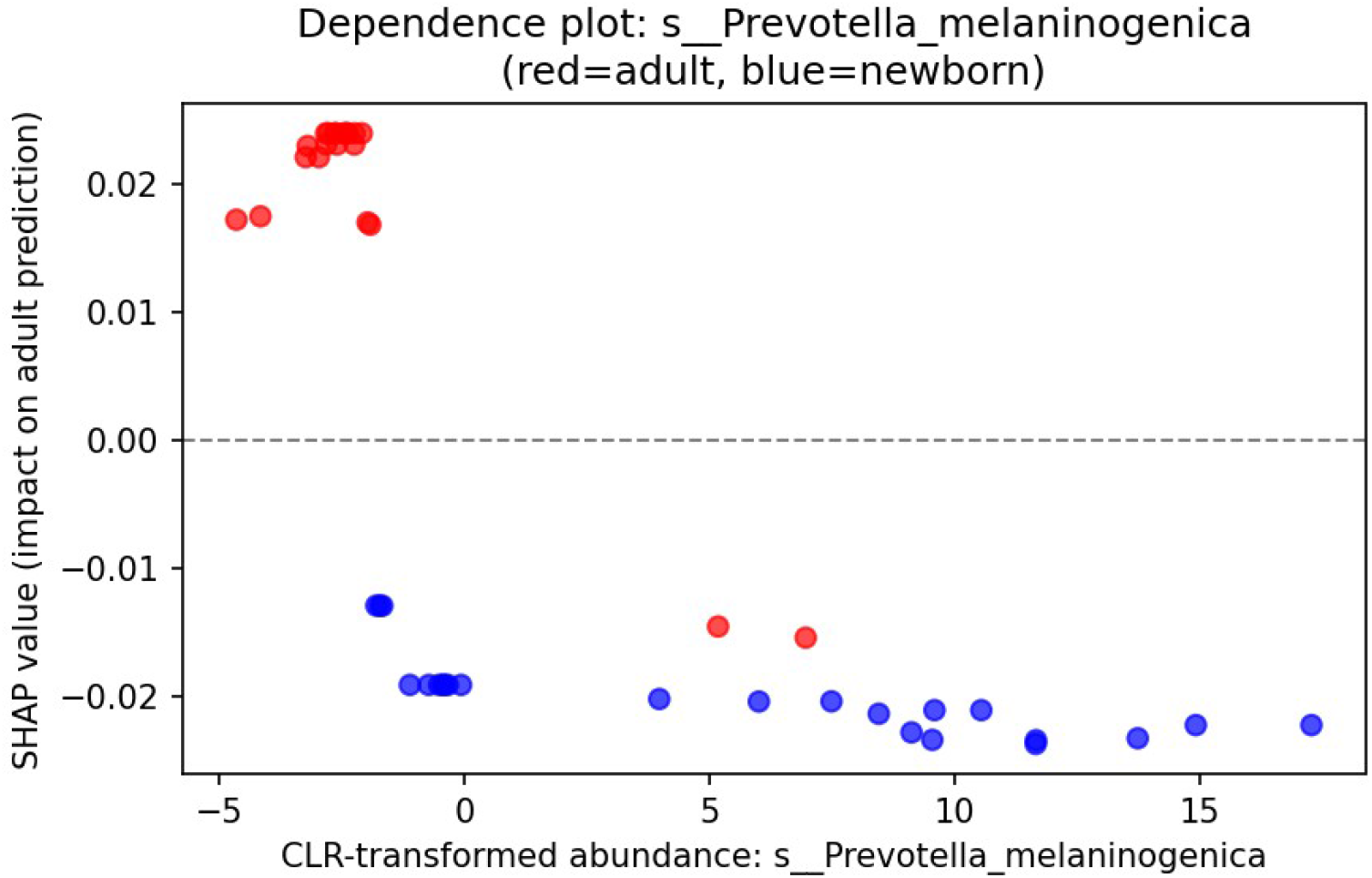
Dependence plots for the three most literature-ambiguous taxa. All three show a clean, strongly negative abundance-SHAP relationship, confirming higher abundance consistently indicates newborn status.

## Discussion

This study set out to answer a methodological question and a biological question together, across two independent cohorts. Key findings:

- **Gut microbiome data alone achieves near-perfect classification** of adult versus newborn status (AUC = 1.00), a ceiling confirmed as genuine signal through a null baseline, bootstrap confidence intervals, and preprocessing sensitivity checks.
- **The fused model nonetheless relies substantially on oral cavity information** (58.1% of SHAP importance versus 41.9% for stool in the primary cohort), demonstrating that a flat accuracy curve can conceal real, non-redundant reliance on a second data source.
- **This site-attribution asymmetry replicates in an independent, larger cohort** (Brito et al., 2016; n=140, sex classification), where oral cavity again contributes more SHAP importance (60.7%/39.3%), and the asymmetry is statistically significant under permutation testing (p < 0.005). The non-significance in the smaller primary cohort (p = 0.89) is consistent with limited statistical power, not absence of the effect.
- **The dominant site flips between cohorts by accuracy but not by SHAP**. In the primary cohort, stool achieves the best single-site AUC; in the Brito cohort, oral cavity does. Yet in both cohorts, the fused model allocates more SHAP importance to oral cavity. This dissociation between accuracy-based and interpretability-based site rankings is itself a novel finding.
- **The taxa driving predictions behave consistently with established microbial colonization biology** once checked directly against per-subject abundance data.
- **An independent cohort study (Myers et al**., **2025)** using 242 paired oral-stool subjects and a different method reports a compatible pattern of complementary, non-redundant signal between the two sites.

### Methodological implications

The accuracy ceiling observed in the primary cohort is unsurprising: adult and newborn microbiomes are among the most compositionally distinct biological states one can compare. The Brito replication cohort deliberately used a harder target (sex-based microbiome differences), and even there, fusion produced no accuracy improvement over the best single site. What is consistent across both cohorts is the SHAP site-attribution pattern: the fused model draws substantially on both sites regardless of whether the accuracy metric can detect this. This cross-cohort consistency strengthens the claim that the SHAP site-attribution framework captures a real, task-independent feature of oral-gut microbiome information content. We have implemented this method as a reusable, publicly available Python tool (available at https://github.com/Aseer47/oral-gut-microbiome-fusion), enabling others to apply the same framework to their own paired multi-site datasets.

### Biological implications

The complementary-signal finding carries a consistent biological reading across both cohorts. If oral and gut microbiomes were maturing or varying in lockstep, a fused model would find little reason to draw heavily on oral information once gut information already solved the task. That both cohorts show substantial oral SHAP importance suggests that the two habitats encode genuinely different biological information about the host. This is consistent with Myers et al.’s finding that over 90% of taxa found in one habitat are absent from the other. A practical implication for microbiome research design: studies aimed at understanding host biological state may systematically underestimate the total available microbiome signal by sampling only the gut.

### Limitations

This study is limited by modest primary cohort sample size (n=44), which produced a directionally consistent but statistically non-significant permutation test result. The replication cohort (n=140) directly addresses this limitation and produces a statistically significant result, but the two cohorts use different biological contrasts (age-group vs. sex), so the replication is of the methodological pattern, not of a single specific biological finding. The dependence analysis was conducted only for the primary cohort’s top-ranked taxa. All findings use Random Forest as the primary model; whether other model families produce similar site-attribution patterns is an open question. Both cohorts are processed with the same MetaPhlAn3 pipeline; external validation using different processing pipelines would further strengthen generalizability claims.

## Conclusion

Combining gut and oral microbiome data from the same subjects did not improve classification accuracy beyond what the best single site achieves in either cohort studied here. That said, SHAP site-attribution consistently showed substantial reliance on oral cavity features in fused models from both cohorts (58.1% and 60.7% respectively), and a permutation test confirmed this asymmetry is statistically significant at the larger sample size. The specific microbial species driving these patterns behave exactly as established early-life colonization biology predicts, once checked empirically rather than assumed from literature. Together, these results make the same underlying point: understanding whether and how multiple body sites relate to each other requires looking past accuracy scores and into what a model actually does with the data it is given.

## Statements and Declarations

### Author Contributions

C.A.R.: Methodology, Software, Formal Analysis, Data Curation, Visualization, Writing – Review & Editing. T.H.S.: Methodology, Software, Formal Analysis, Data Curation, Visualization, Writing – Review & Editing. N.E.S.J.: Conceptualization, Writing – Original Draft, Writing – Review & Editing. A.N.M.T.A.: Conceptualization, Writing – Original Draft, Writing – Review & Editing. C.A.R. and T.H.S. contributed equally as co-first authors. N.E.S.J. and A.N.M.T.A. contributed equally as co-second authors. All authors reviewed and approved the final manuscript.

### Author approvals

All authors have seen and approved the final version of this manuscript, and confirm that it has not been published or accepted for publication elsewhere.

### Data availability

The datasets analyzed in this study are publicly available through the curatedMetagenomicData Bioconductor package (Ferretti et al., 2018 and Brito et al., 2016 cohorts). All analysis outputs, including result tables, figures, complete analysis notebooks, and the standalone site-attribution Python tool, are available at https://github.com/Aseer47/oral-gut-microbiome-fusion.

### Ethics approval

Not applicable. This study used only publicly available, de-identified microbiome sequencing data and did not involve new human subject recruitment or sample collection by the authors.

